## Supplementary material for "Dynamics of genetic code evolution: The emergence of universality": compressed file with latex and all figures: references.bib

@article {nigel,
	author = {Vetsigian, Kalin and Woese, Carl and Goldenfeld, Nigel},
	title = {Collective evolution and the genetic code},
	volume = {103},
	number = {28},
	pages = {10696--10701},
	year = {2006},
	doi = {10.1073/pnas.0603780103},
	publisher = {National Academy of Sciences},
	abstract = {A dynamical theory for the evolution of the genetic code is presented, which accounts for its universality and optimality. The central concept is that a variety of collective, but non-Darwinian, mechanisms likely to be present in early communal life generically lead to refinement and selection of innovation-sharing protocols, such as the genetic code. Our proposal is illustrated by using a simplified computer model and placed within the context of a sequence of transitions that early life may have made, before the emergence of vertical descent.},
	issn = {0027-8424},
	URL = {https://www.pnas.org/content/103/28/10696},
	eprint = {https://www.pnas.org/content/103/28/10696.full.pdf},
	journal = {Proceedings of the National Academy of Sciences}
}
@Article{Sella,
author="Sella, Guy 
and Ardell, David H. ",
title="The Impact of Message Mutation on the Fitness of a Genetic Code",
journal="Journal of Molecular Evolution",
year="2002",
month="May",
day="01",
volume="54",
number="5",
pages="638--651",
abstract="The Standard Genetic Code is organized such that similar codons encode similar amino acids. One explanation suggested that the Standard Code is the result of natural selection to reduce the fitness ``load'' that derives from the mutation and mistranslation of protein-coding genes. We review the arguments against the mutational load-minimizing hypothesis and argue that they need to be reassessed. We review recent analyses of the organization of the Standard Code and conclude that under cautious interpretation they support the mutational load-minimizing hypothesis. We then present a deterministic asexual model with which we study the mode of selection for load minimization. In this model, individual fitness is determined by a protein phenotype resulting from the translation of a mutable set of protein-coding genes. We show that an equilibrium fitness may be associated with a population with the same genetic code and that genetic codes that assign similar codons to similar amino acids have a higher fitness. We also show that the number of mutant codons in each individual at equilibrium, which determines the strength of selection for load minimization, reflects a long-term evolutionary balance between mutations in messages and selection on proteins, rather than the number of mutations that occur in a single generation, as has been assumed by previous authors. We thereby establish that selection for mutational load minimization acts at the level of an individual in a single generation. We conclude with comments on the shortcomings and advantages of load minimization over other hypotheses for the origin of the Standard Code.",
issn="1432-1432",
doi="10.1007/s00239-001-0060-7",
url="https://doi.org/10.1007/s00239-001-0060-7"
}
@MISC{vestigian,
    author = {Kalin Horen Vetsigian},
    title = {COLLECTIVE EVOLUTION OF BIOLOGICAL AND PHYSICAL SYSTEMS},
    year = {2005},
    url="http://guava.physics.uiuc.edu/people/Theses/Vetsigian_PhD_thesis.pdf"
}
@article{goldenfeld,
author = {Butler, Thomas and Goldenfeld, Nigel and Mathew, Damien and Luthey Schulten, Zaida},
year = {2009},
month = {07},
pages = {060901},
title = {Extreme genetic code optimality from a molecular dynamics calculation of amino acid polar requirement},
volume = {79},
journal = {Physical review. E, Statistical, nonlinear, and soft matter physics},
doi = {10.1103/PhysRevE.79.060901}
}
@book{Green,
      author         = "Green, Michael B. and Schwarz, J. H. and Witten, Edward",
      title          = "{SUPERSTRING THEORY. VOL. 2: LOOP AMPLITUDES, ANOMALIES
                        AND PHENOMENOLOGY}",
      year           = "1988",
      url            = "http://www.cambridge.org/us/academic/subjects/physics/theoretical-physics-and-mathematical-physics/superstring-theory-volume-2",
      journal        = "Cambridge, Uk: Univ. Pr. ( 1987) 596 P. ( Cambridge
                        Monographs On Mathematical Physics)",
      ISBN           = "9780521357531",
      SLACcitation   = "%%CITATION = INSPIRE-252419;%%"
}
@book{Alberts,
  title={Molecular Biology of the Cell},
  author={Alberts, B.},
  isbn={9781317563754},
  url={https://books.google.co.uk/books?id=jK6UBQAAQBAJ},
  year={2017},
  publisher={CRC Press}
}
@Article{Sella2006,
author="Sella, Guy
and Ardell, David H.",
title="The Coevolution of Genes and Genetic Codes: Crick's Frozen Accident Revisited",
journal="Journal of Molecular Evolution",
year="2006",
month="Sep",
day="01",
volume="63",
number="3",
pages="297--313",
abstract="The standard genetic code is the nearly universal system for the translation of genes into proteins. The code exhibits two salient structural characteristics: it possesses a distinct organization that makes it extremely robust to errors in replication and translation, and it is highly redundant. The origin of these properties has intrigued researchers since the code was first discovered. One suggestion, which is the subject of this review, is that the code's organization is the outcome of the coevolution of genes and genetic codes. In 1968, Francis Crick explored the possible implications of coevolution at different stages of code evolution. Although he argues that coevolution was likely to influence the evolution of the code, he concludes that it falls short of explaining the organization of the code we see today. The recent application of mathematical modeling to study the effects of errors on the course of coevolution, suggests a different conclusion. It shows that coevolution readily generates genetic codes that are highly redundant and similar in their error-correcting organization to the standard code. We review this recent work and suggest that further affirmation of the role of coevolution can be attained by investigating the extent to which the outcome of coevolution is robust to other influences that were present during the evolution of the code.",
issn="1432-1432",
doi="10.1007/s00239-004-0176-7",
url="https://doi.org/10.1007/s00239-004-0176-7"
}
@article{exception,
author = {Bonitz, S.G. and Berlani, Roberta and Coruzzi, Gloria and Li, May and Macino, Giuseppe and Nobrega, F.G. and Nobrega, M.P. and Thalenfeld, B.E. and Tzagoloff, Alexander},
year = {1980},
month = {07},
pages = {3167-70},
title = {Codon recognition rules in yeast mitochondria},
volume = {77},
journal = {Proceedings of the National Academy of Sciences of the United States of America},
doi = {10.1073/pnas.77.6.3167}
}
@article{1inamillion,
author = {J. Freeland, Stephen and D. Hurst, Laurence},
year = {1998},
month = {10},
pages = {238-48},
title = {The Genetic Code Is One in a Million},
volume = {47},
journal = {Journal of molecular evolution},
doi = {10.1007/PL00006381}
}
@article{woese,
author = {Woese, C.R. and Dugre, D.H. and Saxinger, Carl and Dugre, S.A.},
year = {1966},
month = {05},
pages = {966-74},
title = {The Molecular Basis for the Genetic Code},
volume = {55},
journal = {Proceedings of the National Academy of Sciences of the United States of America},
doi = {10.1073/pnas.55.4.966}
}
@Article{Mathew2008,
author="Mathew, Damien C.
and Luthey-Schulten, Zaida",
title="On the Physical Basis of the Amino Acid Polar Requirement",
journal="Journal of Molecular Evolution",
year="2008",
month="May",
day="01",
volume="66",
number="5",
pages="519--528",
abstract="Understanding how codons became associated with their specific amino acids is fundamental to deriving a theory for the origin of the genetic code. Carl Woese and coworkers designed a series of experiments to test associations between amino acids and nucleobases that may have played a role in establishing the genetic code. Through these experiments it was found that a property of amino acids called the polar requirement (PR) is correlated with the organization of the codon table. No other property of amino acids has been found that correlates with the codon table as well as PR, indicating that PR is uniquely related to the modern genetic code. Using molecular dynamics simulations of amino acids in solutions of water and dimethylpyridine used to experimentally measure PR, we show that variations in the partitioning between the two phases as described by radial distribution functions correlate well with the measured PRs. Partition coefficients based on probability densities of the amino acids in each phase have the linear behavior with base concentration as suggested by PR experiments.",
issn="1432-1432",
doi="10.1007/s00239-008-9073-9",
url="https://doi.org/10.1007/s00239-008-9073-9"
}
@article{butler,
author = {Butler, Thomas and Goldenfeld, Nigel and Mathew, Damien and Luthey Schulten, Zaida},
year = {2009},
month = {07},
pages = {060901},
title = {Extreme genetic code optimality from a molecular dynamics calculation of amino acid polar requirement},
volume = {79},
journal = {Physical review. E, Statistical, nonlinear, and soft matter physics},
doi = {10.1103/PhysRevE.79.060901}
}
@Article{Freeland2003,
author="Freeland, Stephen J.
and Wu, Tao
and Keulmann, Nick",
title="The Case for an Error Minimizing Standard Genetic Code",
journal="Origins of life and evolution of the biosphere",
year="2003",
month="Oct",
day="01",
volume="33",
number="4",
pages="457--477",
abstract="Since discovering the pattern by which amino acids are assigned to codons within the standard genetic code, investigators have explored the idea that natural selection placed biochemically similar amino acids near to one another in coding space so as to minimize the impact of mutations and/or mistranslations. The analytical evidence to support this theory has grown in sophistication and strength over the years, and counterclaims questioning its plausibility and quantitative support have yet to transcend some significant weaknesses in their approach. These weaknesses are illustrated here by means of a simple simulation model for adaptive genetic code evolution. There remain ill explored facets of the `error minimizing' code hypothesis, however, including the mechanism and pathway by which an adaptive pattern of codon assignments emerged, the extent to which natural selection created synonym redundancy, its role in shaping the amino acid and nucleotide languages, and even the correct interpretation of the adaptive codon assignment pattern: these represent fertile areas for future research.",
issn="1573-0875",
doi="10.1023/A:1025771327614",
url="https://doi.org/10.1023/A:1025771327614"
}
@article{ardell,
author = {Becich, Peter and P Stark, Brian and S Bhat, Harish and Ardell, David},
year = {2013},
month = {02},
pages = {111-25},
title = {CMCpy: Genetic Code-Message Coevolution Models in Python},
volume = {9},
journal = {Evolutionary bioinformatics online},
doi = {10.4137/EBO.S11169}
}
@article {Woese8742,
	author = {Woese, Carl R.},
	title = {On the evolution of cells},
	volume = {99},
	number = {13},
	pages = {8742--8747},
	year = {2002},
	doi = {10.1073/pnas.132266999},
	publisher = {National Academy of Sciences},
	abstract = {A theory for the evolution of cellular organization is presented. The model is based on the (data supported) conjecture that the dynamic of horizontal gene transfer (HGT) is primarily determined by the organization of the recipient cell. Aboriginal cell designs are taken to be simple and loosely organized enough that all cellular componentry can be altered and/or displaced through HGT, making HGT the principal driving force in early cellular evolution. Primitive cells did not carry a stable organismal genealogical trace. Primitive cellular evolution is basically communal. The high level of novelty required to evolve cell designs is a product of communal invention, of the universal HGT field, not intralineage variation. It is the community as a whole, the ecosystem, which evolves. The individual cell designs that evolved in this way are nevertheless fundamentally distinct, because the initial conditions in each case are somewhat different. As a cell design becomes more complex and interconnected a critical point is reached where a more integrated cellular organization emerges, and vertically generated novelty can and does assume greater importance. This critical point is called the {\textquotedblleft}Darwinian Threshold{\textquotedblright} for the reasons given. HGT,horizontal gene transfer;SMA,supramolecular aggregate},
	issn = {0027-8424},
	URL = {https://www.pnas.org/content/99/13/8742},
	eprint = {https://www.pnas.org/content/99/13/8742.full.pdf},
	journal = {Proceedings of the National Academy of Sciences}
}
@article{Yang,
      author         = "He, Yang-Hui",
      title          = "{The Calabi-Yau Landscape: from Geometry, to Physics, to
                        Machine-Learning}",
      year           = "2018",
      eprint         = "1812.02893",
      url="https://arxiv.org/abs/1812.02893",
      archivePrefix  = "arXiv",
      primaryClass   = "hep-th",
      SLACcitation   = "%%CITATION = ARXIV:1812.02893;%%"
      
}
@Article{Haig1991,
author="Haig, David
and Hurst, Laurence D.",
title="A quantitative measure of error minimization in the genetic code",
journal="Journal of Molecular Evolution",
year="1991",
month="Nov",
day="01",
volume="33",
number="5",
pages="412--417",
abstract="We have calculated the average effect of changing a codon by a single base for all possible single-base changes in the genetic code and for changes in the first, second, and third codon positions separately. Such values were calculated for an amino acid's polar requirement, hydropathy, molecular volume, and isoelectric point. For each attribute the average effect of single-base changes was also calculated for a large number of randomly generated codes that retained the same level of redundancy as the natural code. Amino acids whose codons differed by a single base in the first and third codon positions were very similar with respect to polar requirement and hydropathy. The major differences between amino acids were specified by the second codon position. Codons with U in the second position are hydrophobic, whereas most codons with A in the second position are hydrophilic. This accounts for the observation of complementary hydropathy. Single-base changes in the natural code had a smaller average effect on polar requirement than all but 0.02{\%} of random codes. This result is most easily explained by selection to minimize deleterious effects of translation errors during the early evolution of the code.",
issn="1432-1432",
doi="10.1007/BF02103132",
url="https://doi.org/10.1007/BF02103132"
}
@book{feller2008introduction,
  title={AN INTRODUCTION TO PROBABILITY THEORY AND ITS APPLICATIONS, 2ND ED},
  author={Feller, W.},
  number={v. 2},
  isbn={9788126518067},
  url={https://books.google.co.uk/books?id=OXkg-LvRgjUC},
  year={2008},
  publisher={Wiley India Pvt. Limited}
}
@article{Flajolet1992BirthdayPC,
  title={Birthday Paradox, Coupon Collectors, Caching Algorithms and Self-Organizing Search},
  author={Philippe Flajolet and Dani{\`e}le Gardy and Lo{\"y}s Thimonier},
  journal={Discrete Applied Mathematics},
  year={1992},
  volume={39},
  pages={207-229}
}
@book{baylis2017error,
  title={Error Correcting Codes: A Mathematical Introduction},
  author={Baylis, D.J.},
  isbn={9781351449847},
  url={https://books.google.co.uk/books?id=r0OVtAEACAAJ},
  year={2017},
  publisher={Taylor \& Francis}
}
@book{alon2019introduction,
  title={An Introduction to Systems Biology},
  author={Alon, U.},
  isbn={9781439837177},
  lccn={2019003924},
  series={Chapman \& Hall/Crc Mathematical and Computational Biology},
  url={https://books.google.co.uk/books?id=MWXdQgAACAAJ},
  year={2019},
  publisher={CRC Press LLC}
}
@book{Nakahara:2003nw,
      author         = "Nakahara, M.",
      title          = "{Geometry, topology and physics}",
      journal        = "Boca Raton, USA: Taylor & Francis (2003) 573 p",
      year           = "2003",
      SLACcitation   = "%%CITATION = INSPIRE-640779;%%"
}
@article{10.1093/emph/eov018,
    author = {Burmeister, Alita R.},
    title = "{Horizontal Gene Transfer}",
    journal = {Evolution, Medicine, and Public Health},
    volume = {2015},
    number = {1},
    pages = {193-194},
    year = {2015},
    month = {07},
    issn = {2050-6201},
    doi = {10.1093/emph/eov018},
    url = {https://doi.org/10.1093/emph/eov018},
    eprint = {http://oup.prod.sis.lan/emph/article-pdf/2015/1/193/17926586/eov018.pdf},
}
@book{cohen1997covering,
  title={Covering Codes},
  author={Cohen, G. and Honkala, I. and Litsyn, S. and Lobstein, A.},
  isbn={9780080530079},
  lccn={97002461},
  series={North-Holland Mathematical Library},
  url={https://books.google.co.uk/books?id=7KBYOt44sugC},
  year={1997},
  publisher={Elsevier Science}
}
@book{Allenby:2010:CIC:1942905,
 author = {Allenby, R. B.J.T. and Slomson, Alan},
 title = {How to Count: An Introduction to Combinatorics, Second Edition},
 year = {2010},
 isbn = {1420082604, 9781420082609},
 edition = {2nd},
 publisher = {Chapman \& Hall/CRC},
}


@article{Adams:2019kby,
      author         = "Adams, Fred C.",
      title          = "{The Degree of Fine-Tuning in our Universe -- and
                        Others}",
      year           = "2019",
      eprint         = "1902.03928",
      url="https://arxiv.org/abs/1902.03928#",
      archivePrefix  = "arXiv",
      primaryClass   = "astro-ph.CO",
      SLACcitation   = "%%CITATION = ARXIV:1902.03928;%%"
}
@book{strogatz2000nonlinear,
  title={Nonlinear Dynamics and Chaos: With Applications to Physics, Biology, Chemistry and Engineering},
  author={Strogatz, S.H.},
  series={Studies in nonlinearity},
  url={https://books.google.co.uk/books?id=NZZDnQEACAAJ},
  year={2000},
  publisher={Westview}
}
@article {Hoshika884,
	author = {Hoshika, Shuichi and Leal, Nicole A. and Kim, Myong-Jung and Kim, Myong-Sang and Karalkar, Nilesh B. and Kim, Hyo-Joong and Bates, Alison M. and Watkins, Norman E. and SantaLucia, Holly A. and Meyer, Adam J. and DasGupta, Saurja and Piccirilli, Joseph A. and Ellington, Andrew D. and SantaLucia, John and Georgiadis, Millie M. and Benner, Steven A.},
	title = {Hachimoji DNA and RNA: A genetic system with eight building blocks},
	volume = {363},
	number = {6429},
	pages = {884--887},
	year = {2019},
	doi = {10.1126/science.aat0971},
	publisher = {American Association for the Advancement of Science},
	abstract = {DNA and RNA are naturally composed of four nucleotide bases that form hydrogen bonds in order to pair. Hoshika et al. added an additional four synthetic nucleotides to produce an eight-letter genetic code and generate so-called hachimoji DNA. Coupled with an engineered T7 RNA polymerase, this expanded DNA alphabet could be transcribed into RNA. Thus, new forms of DNA that add information density to genetic biopolymers can be generated that may be useful for future synthetic biological applications.Science, this issue p. 884We report DNA- and RNA-like systems built from eight nucleotide {\textquotedblleft}letters{\textquotedblright} (hence the name {\textquotedblleft}hachimoji{\textquotedblright}) that form four orthogonal pairs. These synthetic systems meet the structural requirements needed to support Darwinian evolution, including a polyelectrolyte backbone, predictable thermodynamic stability, and stereoregular building blocks that fit a Schr{\"o}dinger aperiodic crystal. Measured thermodynamic parameters predict the stability of hachimoji duplexes, allowing hachimoji DNA to increase the information density of natural terran DNA. Three crystal structures show that the synthetic building blocks do not perturb the aperiodic crystal seen in the DNA double helix. Hachimoji DNA was then transcribed to give hachimoji RNA in the form of a functioning fluorescent hachimoji aptamer. These results expand the scope of molecular structures that might support life, including life throughout the cosmos.},
	issn = {0036-8075},
	URL = {https://science.sciencemag.org/content/363/6429/884},
	eprint = {https://science.sciencemag.org/content/363/6429/884.full.pdf},
	journal = {Science}
}
@ARTICLE{automorph,
       author = {{Morteza Mirafzal}, S. and {Ziaee}, Meysam},
        title = "{A note on the automorphism group of the Hamming graph}",
      journal = {arXiv e-prints},
     keywords = {Mathematics - Group Theory},
         year = "2019",
        month = "Jan",
          eid = {arXiv:1901.07784},
        pages = {arXiv:1901.07784},
archivePrefix = {arXiv},
       eprint = {1901.07784},
 primaryClass = {math.GR},
       adsurl = {https://ui.adsabs.harvard.edu/abs/2019arXiv190107784M},
      adsnote = {Provided by the SAO/NASA Astrophysics Data System}
}
@book{bialek2012biophysics,
  title={Biophysics: Searching for Principles},
  author={Bialek, W.},
  isbn={9780691138916},
  lccn={2012014443},
  url={https://books.google.co.uk/books?id=5In\_FKA2rmUC},
  year={2012},
  publisher={Princeton University Press}
}
@misc{Petersen2008,
  abstract = {Matrix identities, relations and approximations. A desktop reference for quick overview of mathematics of matrices.},
  added-at = {2011-01-17T12:52:58.000+0100},
  author = {Petersen, K. B. and Pedersen, M. S.},
  biburl = {https://www.bibsonomy.org/bibtex/263c840382cc4b1efb8cefe447465b7ac/hkayabilisim},
  file = {:home/hkaya/Projeler/diagnus/Screener/doc/literature/Petersen2008.pdf:PDF},
  interhash = {6368b9b490c0225e22334ea0a0841a33},
  intrahash = {63c840382cc4b1efb8cefe447465b7ac},
  keywords = {matrixderivative inverse Matrixidentity matrixrelations},
  month = oct,
  note = {Version 20081110},
  publisher = {Technical University of Denmark},
  review = {Matrix Cookbook},
  timestamp = {2011-01-17T12:52:58.000+0100},
  title = {The Matrix Cookbook},
  url = {http://www2.imm.dtu.dk/pubdb/p.php?3274},
  year = 2008
}

@article{Wilson:1974mb,
      author         = "Wilson, Kenneth G.",
      title          = "{The Renormalization Group: Critical Phenomena and the
                        Kondo Problem}",
      journal        = "Rev. Mod. Phys.",
      volume         = "47",
      year           = "1975",
      pages          = "773",
      doi            = "10.1103/RevModPhys.47.773",
      reportNumber   = "CLNS-296",
      SLACcitation   = "%%CITATION = RMPHA,47,773;%%"
}

@article{Wilson:1993dy,
      author         = "Wilson, K. G.",
      title          = "{The renormalization group and critical phenomena}",
      journal        = "Rev. Mod. Phys.",
      volume         = "55",
      year           = "1983",
      pages          = "583-600",
      doi            = "10.1103/RevModPhys.55.583",
      SLACcitation   = "%%CITATION = RMPHA,55,583;%%"
}

@book{Goldenfeld:1992qy,
      author         = "Goldenfeld, N.",
      title          = "{Lectures on phase transitions and the renormalization
                        group}",
      journal        = "Reading, USA: Addison-Wesley (1992) 394 p. (Frontiers in
                        physics, 85)",
      year           = "1992",
      SLACcitation   = "%%CITATION = INSPIRE-345315;%%"
}

@ARTICLE{widom,
   author = {{Widom}, B.},
    title = "{Equation of State in the Neighborhood of the Critical Point}",
  journal = {Journal of Chemical Physics},
     year = 1965,
    month = dec,
   volume = 43,
    pages = {3898-3905},
      doi = {10.1063/1.1696618},
   adsurl = {https://ui.adsabs.harvard.edu/abs/1965JChPh..43.3898W},
  adsnote = {Provided by the SAO/NASA Astrophysics Data System}
}

@article{Denef:2006ad,
      author         = "Denef, Frederik and Douglas, Michael R.",
      title          = "{Computational complexity of the landscape. I.}",
      journal        = "Annals Phys.",
      volume         = "322",
      year           = "2007",
      pages          = "1096-1142",
      doi            = "10.1016/j.aop.2006.07.013",
      eprint         = "hep-th/0602072",
      archivePrefix  = "arXiv",
      primaryClass   = "hep-th",
      SLACcitation   = "%%CITATION = HEP-TH/0602072;%%"
}

@article{Weinberg:1988cp,
      author         = "Weinberg, Steven",
      title          = "{The Cosmological Constant Problem}",
      journal        = "Rev. Mod. Phys.",
      volume         = "61",
      year           = "1989",
      pages          = "1-23",
      doi            = "10.1103/RevModPhys.61.1",
      note           = "[,569(1988)]",
      reportNumber   = "UTTG-12-88",
      SLACcitation   = "%%CITATION = RMPHA,61,1;%%"
}

@inproceedings{Polchinski:2006gy,
      author         = "Polchinski, Joseph",
      title          = "{The Cosmological Constant and the String Landscape}",
      booktitle      = "{The Quantum Structure of Space and Time: Proceedings of
                        the 23rd Solvay Conference on Physics. Brussels, Belgium.
                        1 - 3 December 2005}",
      year           = "2006",
      pages          = "216-236",
      eprint         = "hep-th/0603249",
      archivePrefix  = "arXiv",
      primaryClass   = "hep-th",
      SLACcitation   = "%%CITATION = HEP-TH/0603249;%%"
}

@article{jkm,
      author         = "Jejjala, Vishnu and Kavic, Michael and Minic, Djordje",
      title          = "{Time and M-theory}",
      journal        = "Int. J. Mod. Phys.",
      volume         = "A22",
      year           = "2007",
      pages          = "3317-3405",
      doi            = "10.1142/S0217751X07036981",
      eprint         = "0706.2252",
      archivePrefix  = "arXiv",
      primaryClass   = "hep-th",
      reportNumber   = "DCPT-07-31, VPI-IPNAS-07-05",
      SLACcitation   = "%%CITATION = ARXIV:0706.2252;%%"
}
