## Supplementary figures and images for "Dynamics of genetic code evolution: The emergence of universality"

### 223fitphi001.png

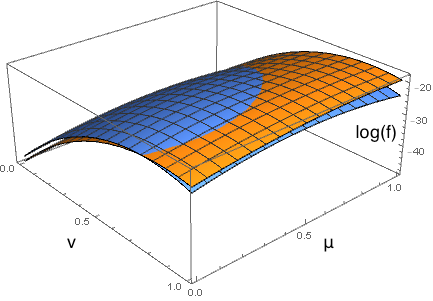

### 223fitphi01.png

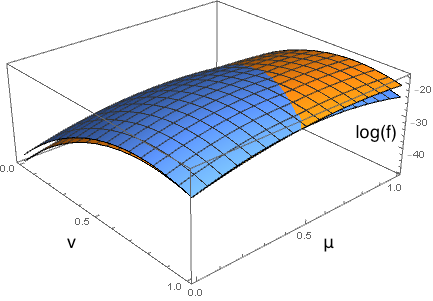

### 223fitphi05.png

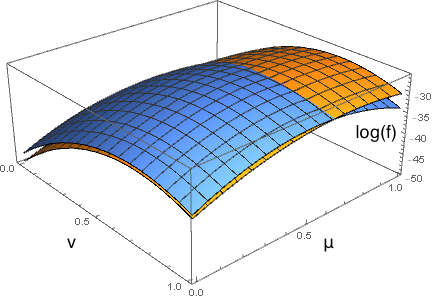

### 223fitphi099.png

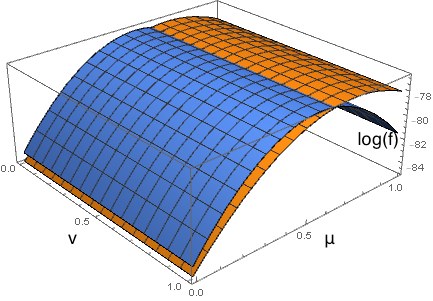

### 223logfphi001.png

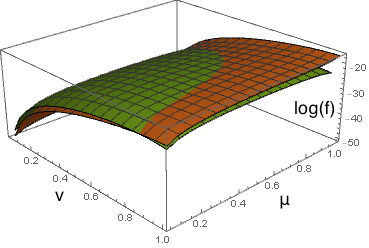

### 223logfphi01.png

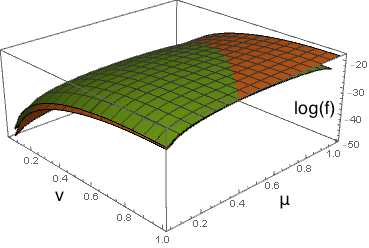

### 223logfphi05.png

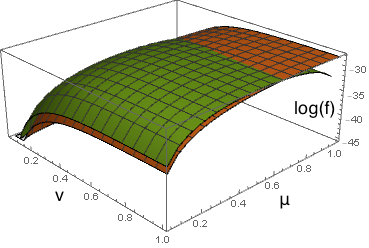

### 223logfphi099.png

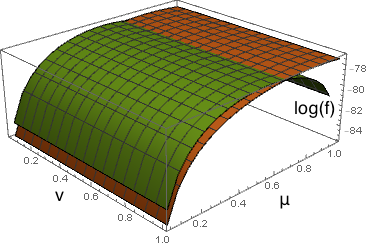

### 413logfphi001.png

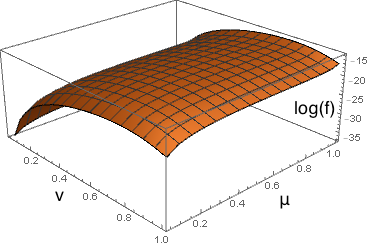

### 413logfphi01.png

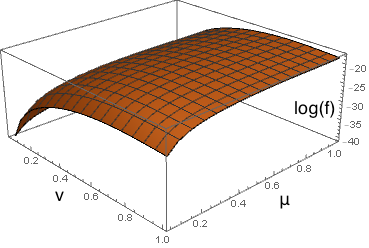

### 413logfphi05.png

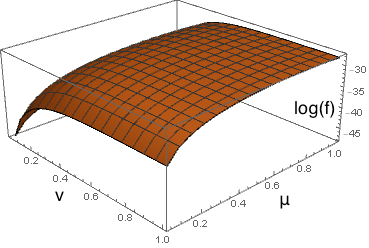

### 413logfphi099.png

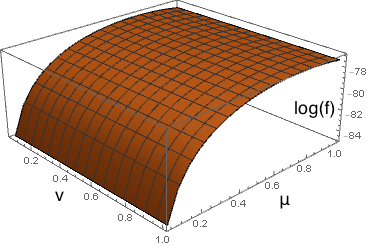

### avaover10.png

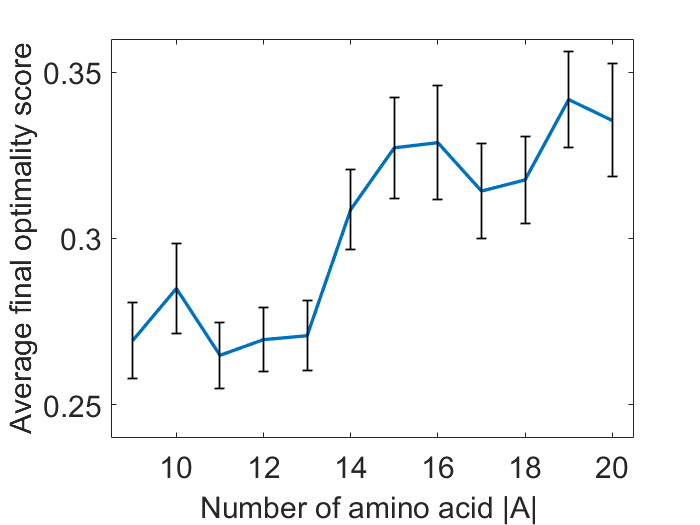

### aventityover10.png

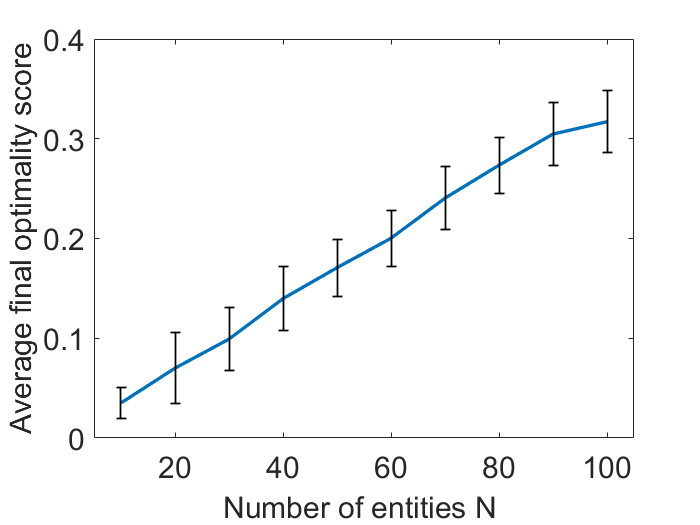

### avHover10.png

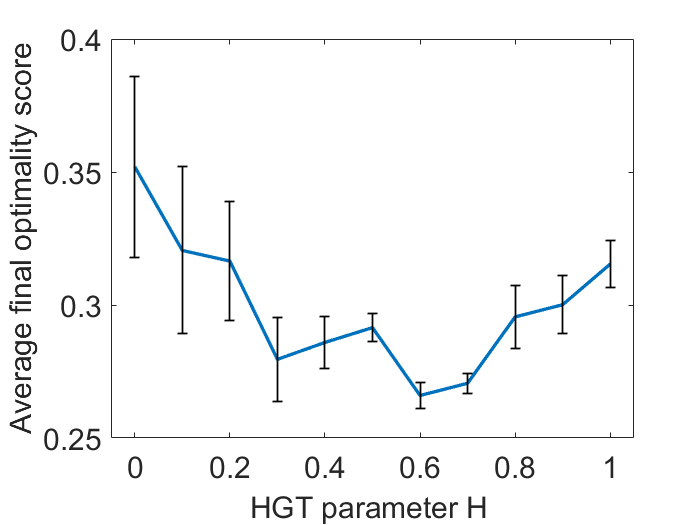

### aviover10.png

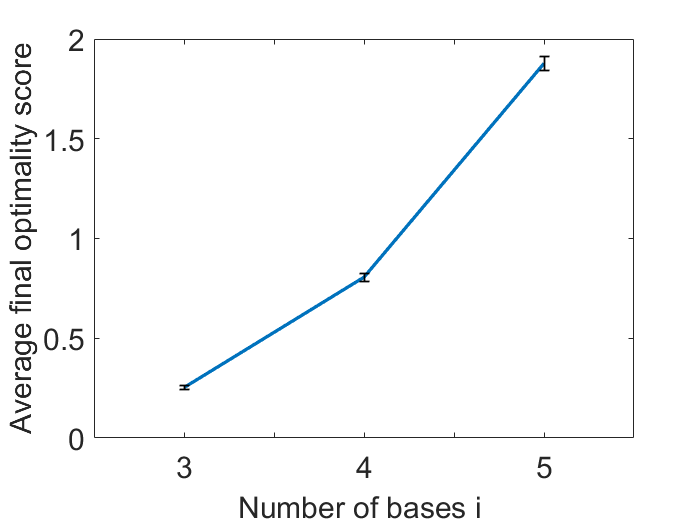

### avKover10.png

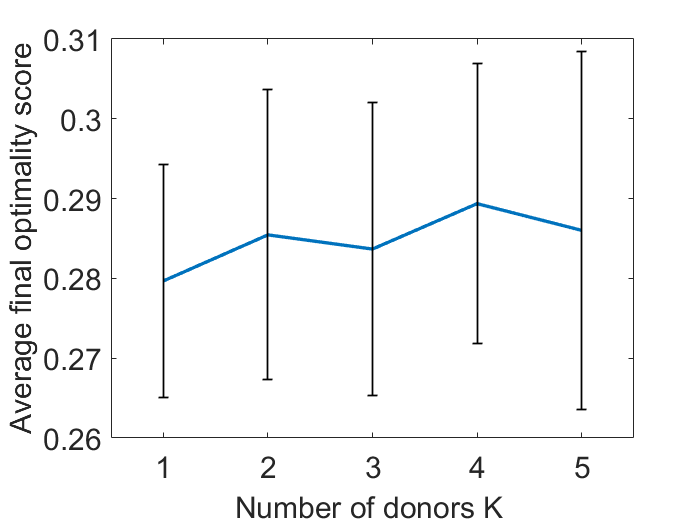

### avmunuover10.png

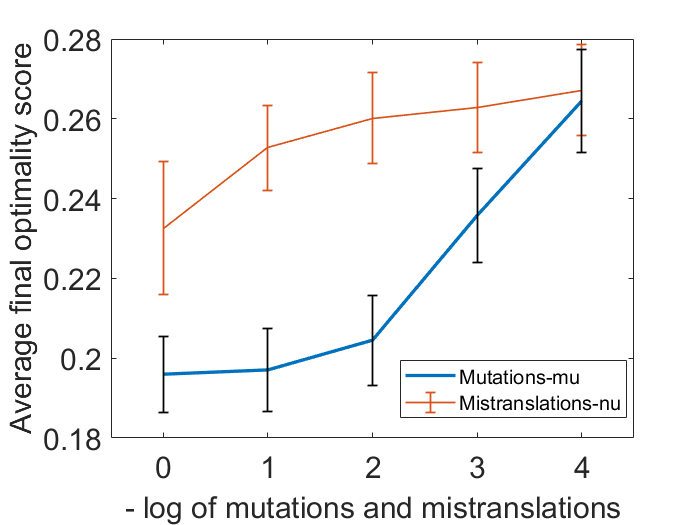

### avnover10.png

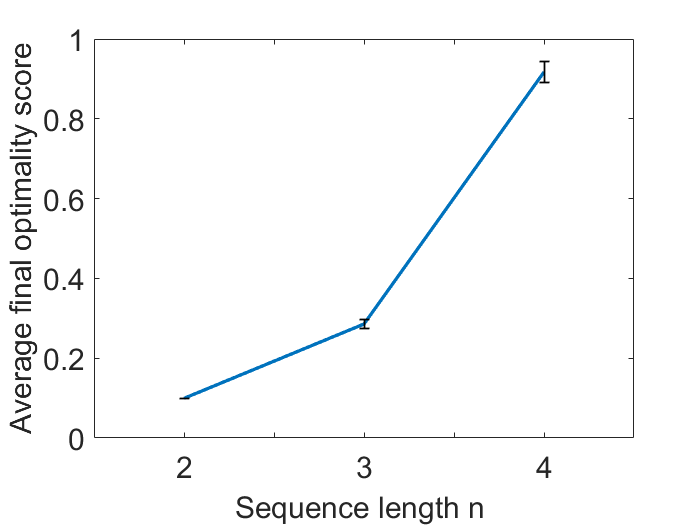

### color_plot_fitness.png

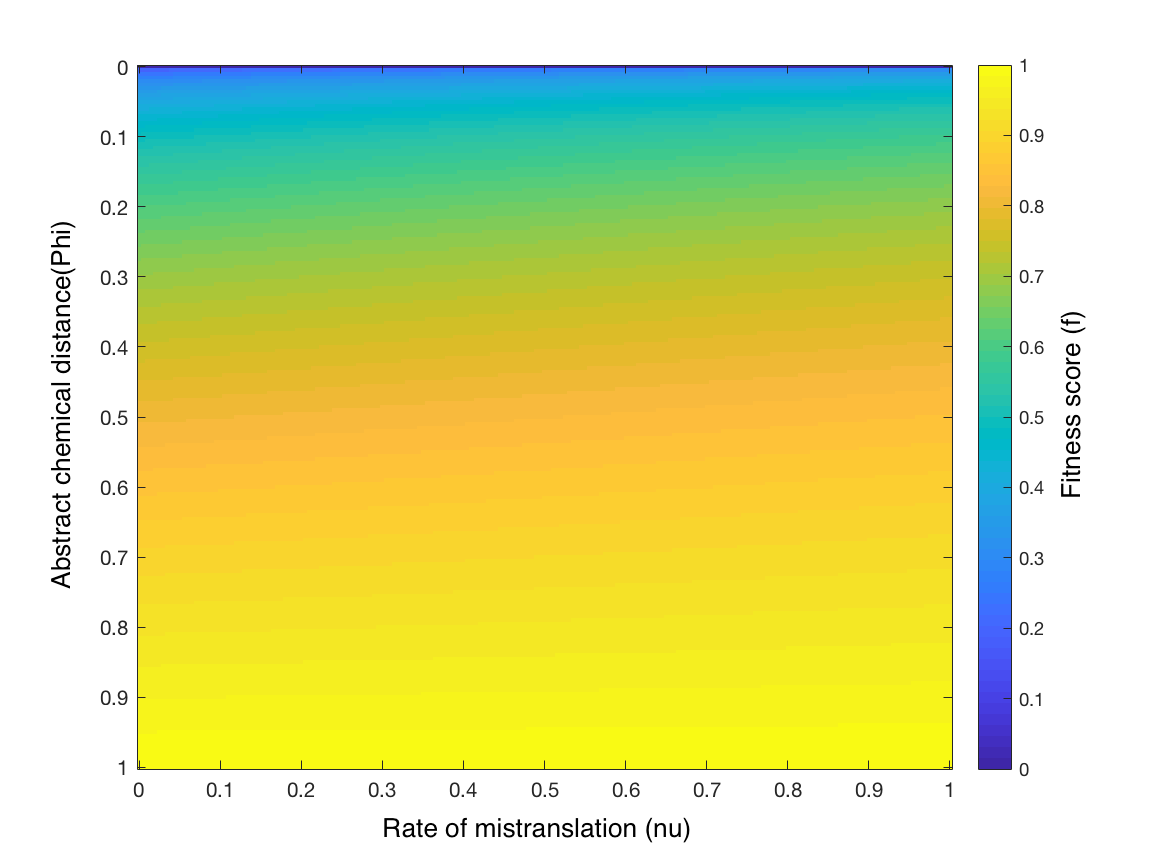

### diffstartresult.png

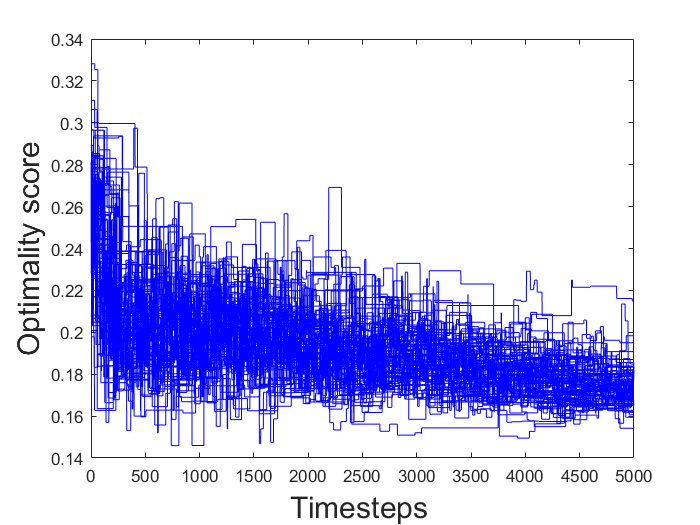

### diffstarttimeevo.png

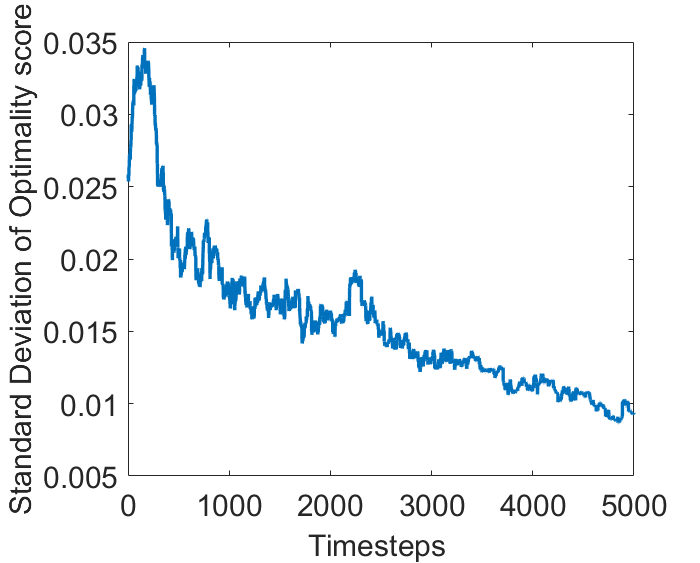

### H0first.png

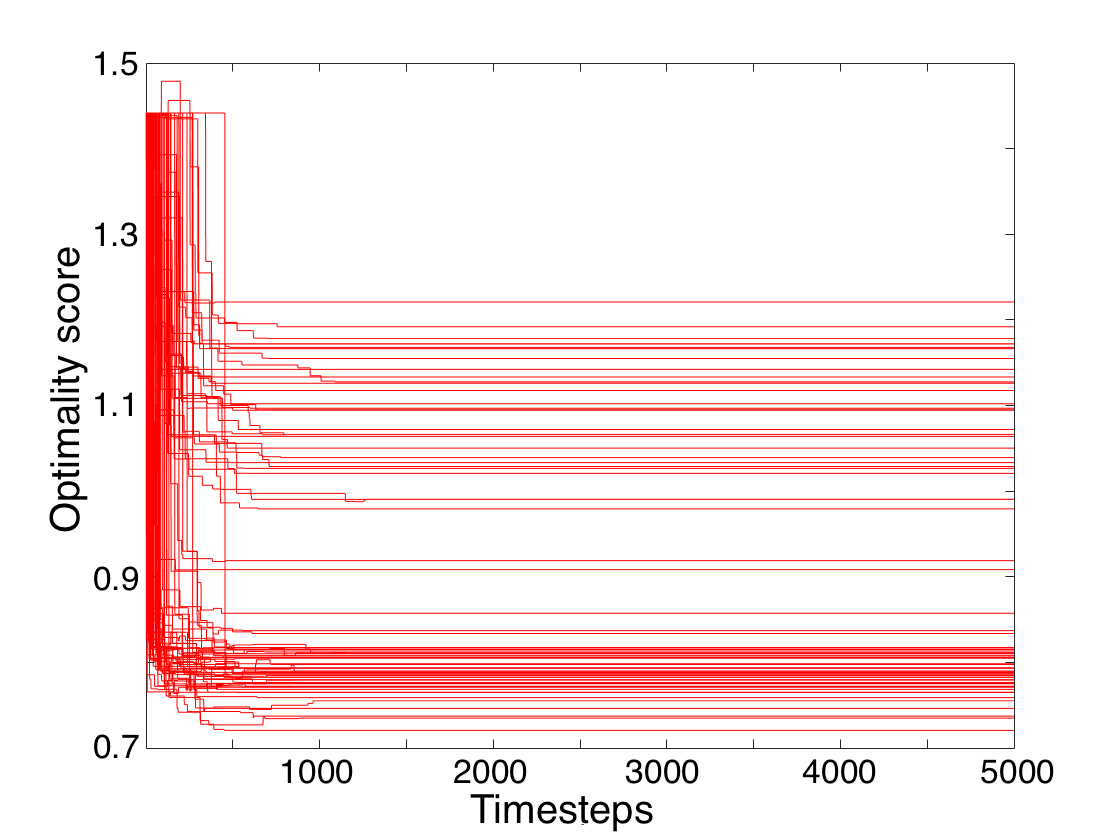

### H04first.png

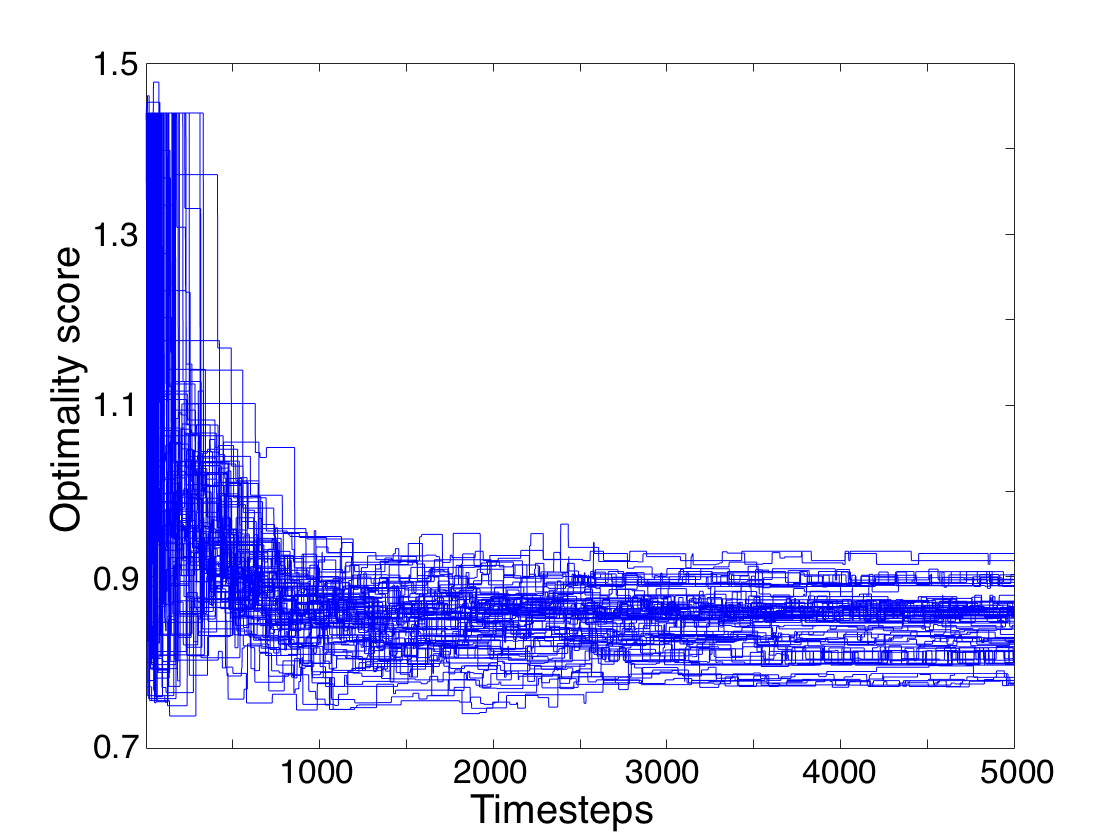

### stdevoa.png

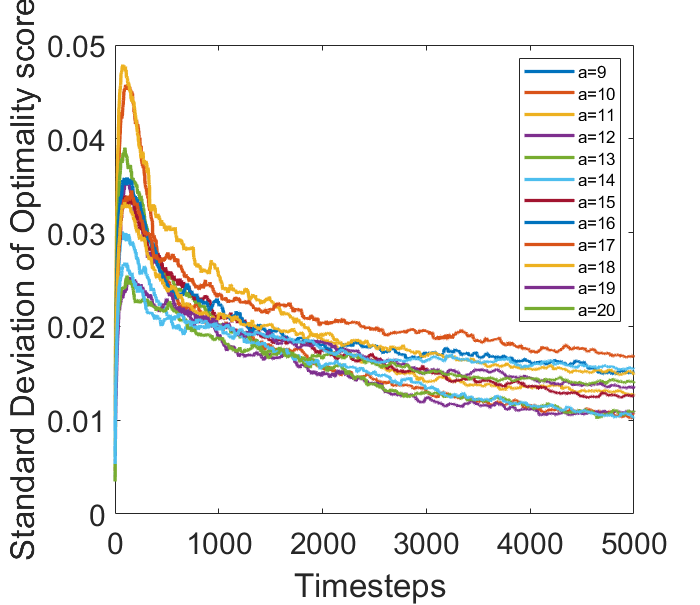

### stdevoentityover10.png

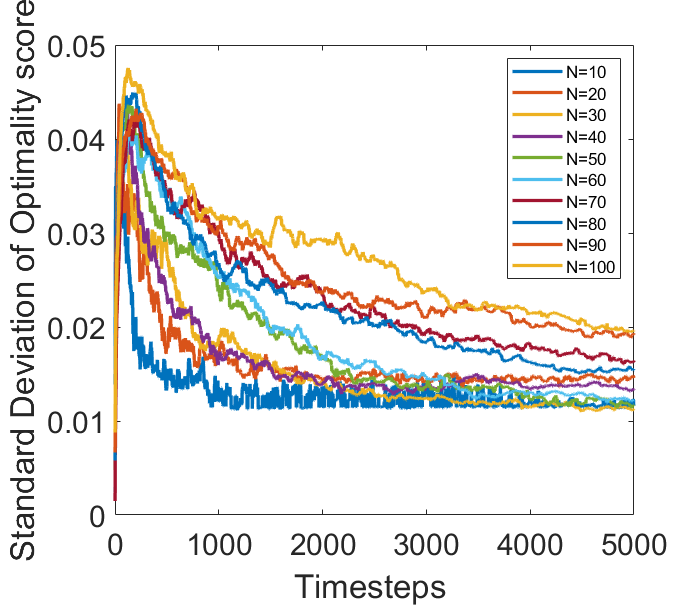

### stdevoH.png

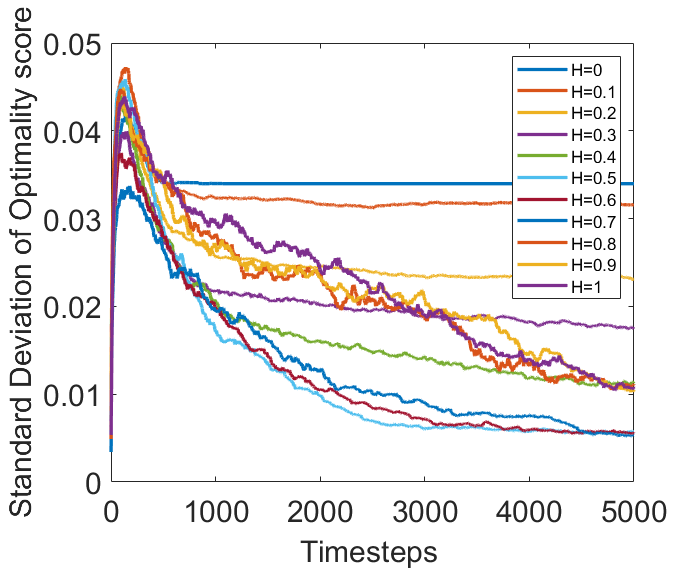

### stdevoi.png

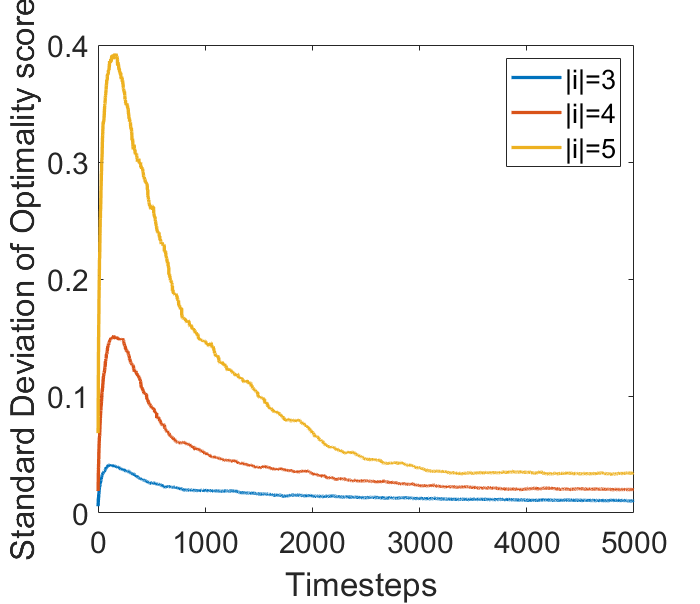

### stdevoK.png

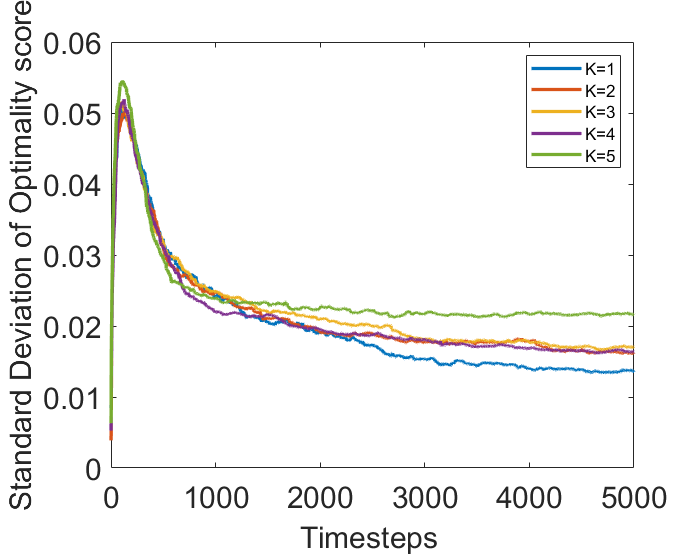

### stdevomunu.png

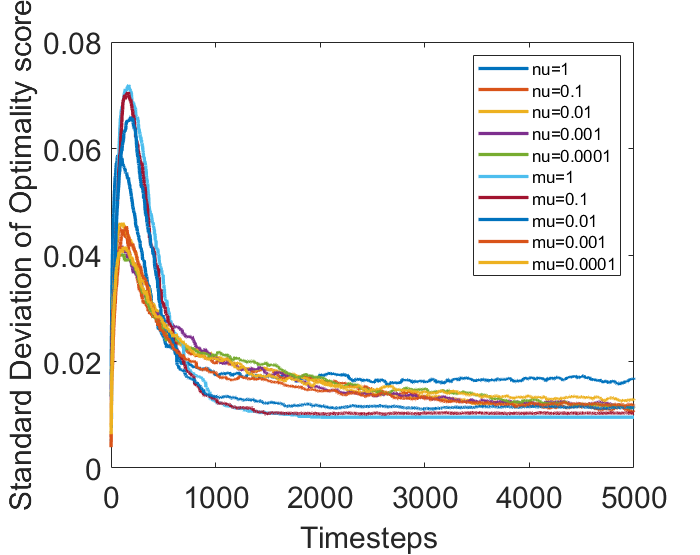

### stdevon.png

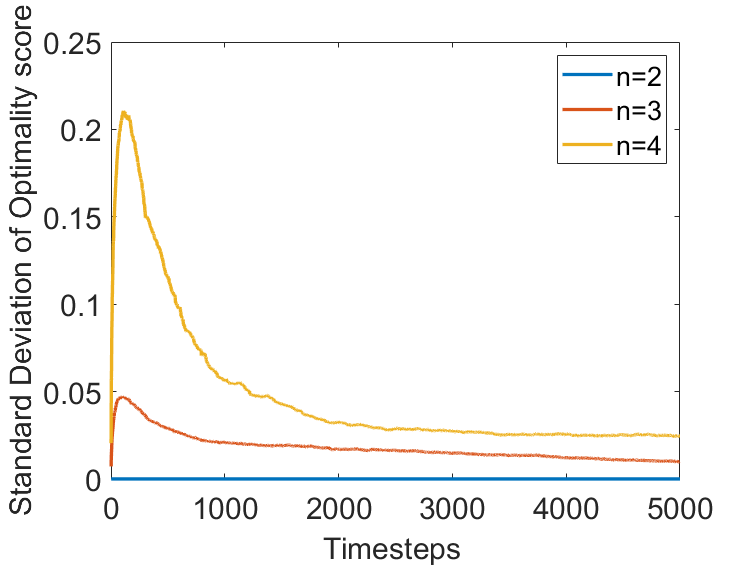

### surface_plot_fitness.png

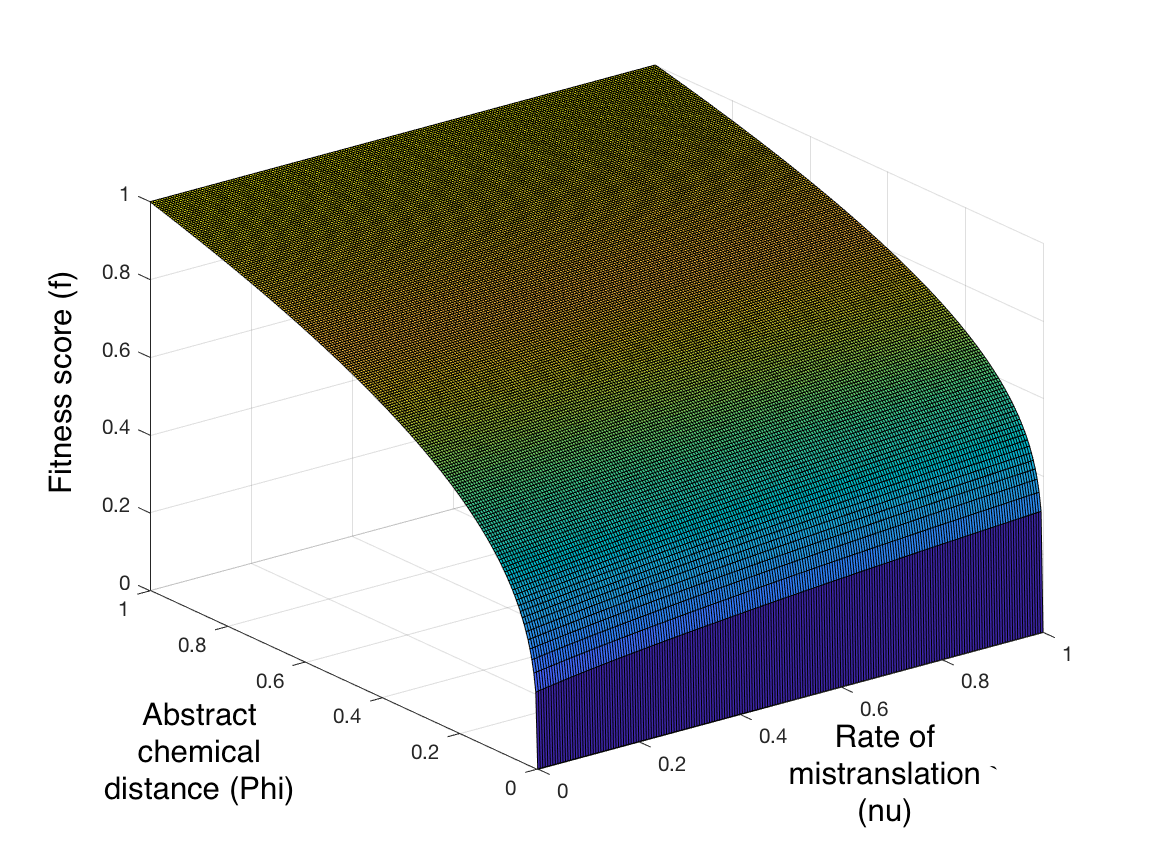

### timevarH5.png

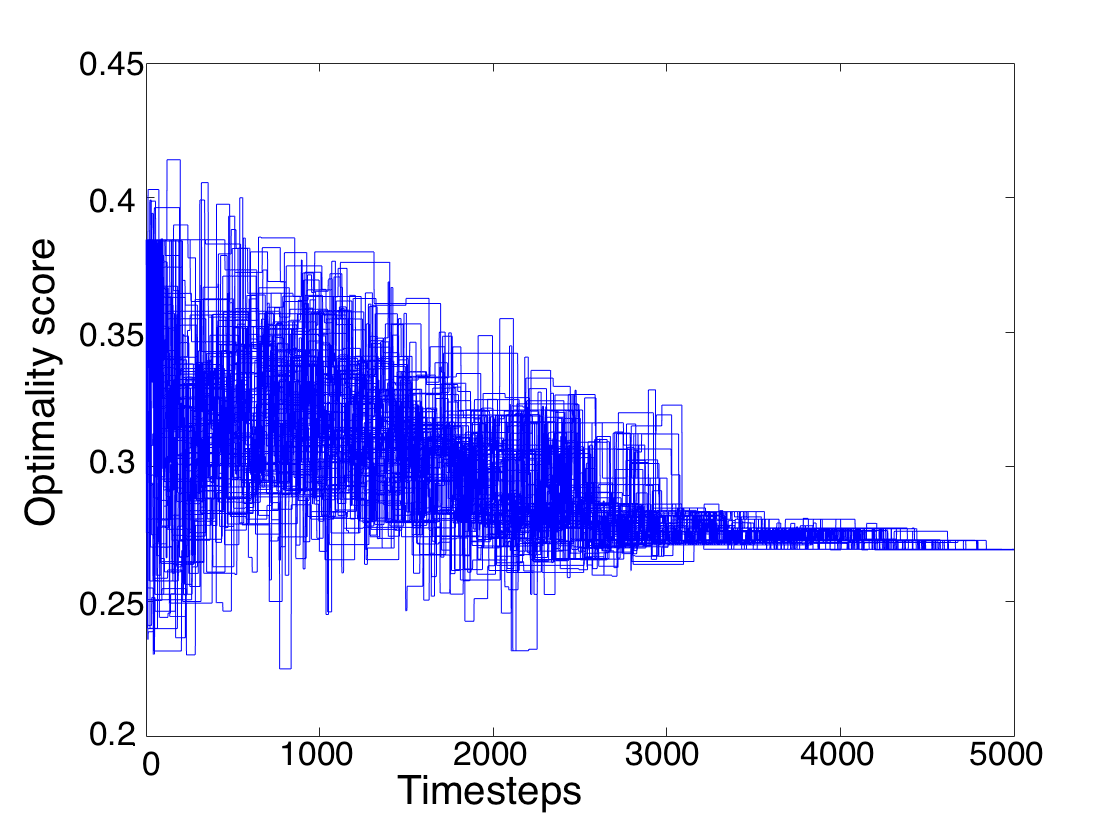

### timevarH6.png

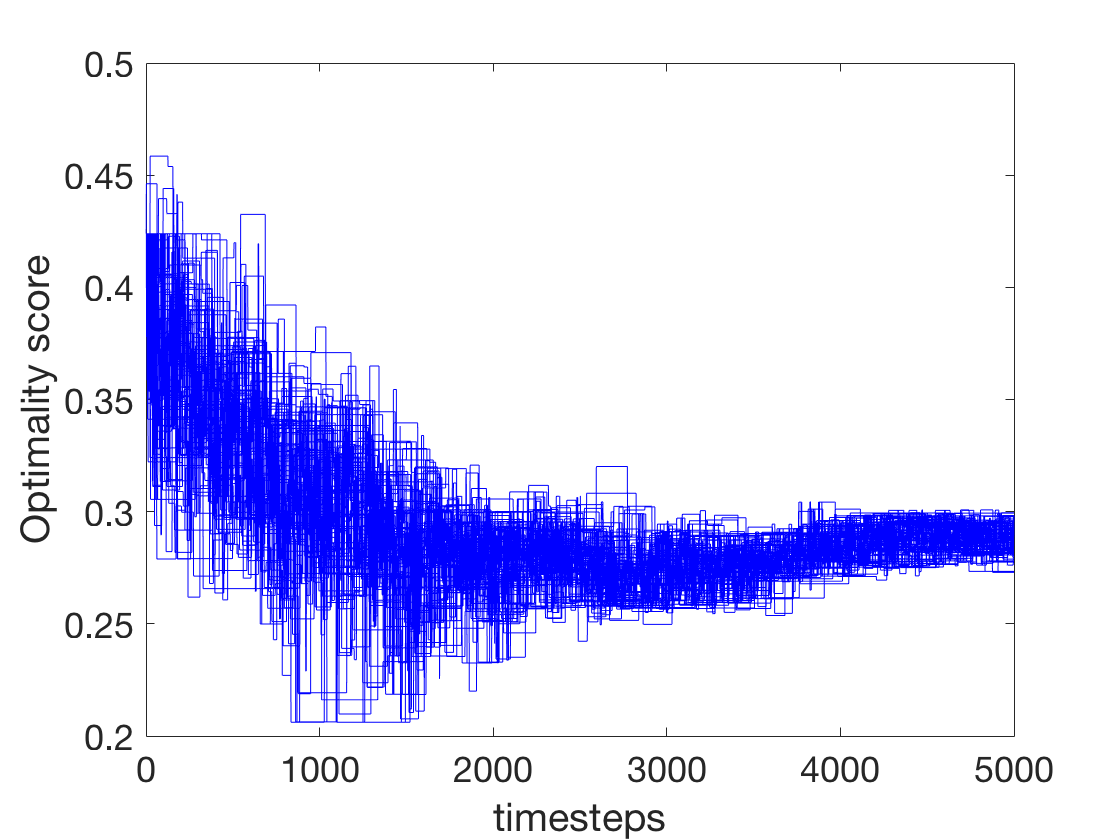

### timevarinH1.png

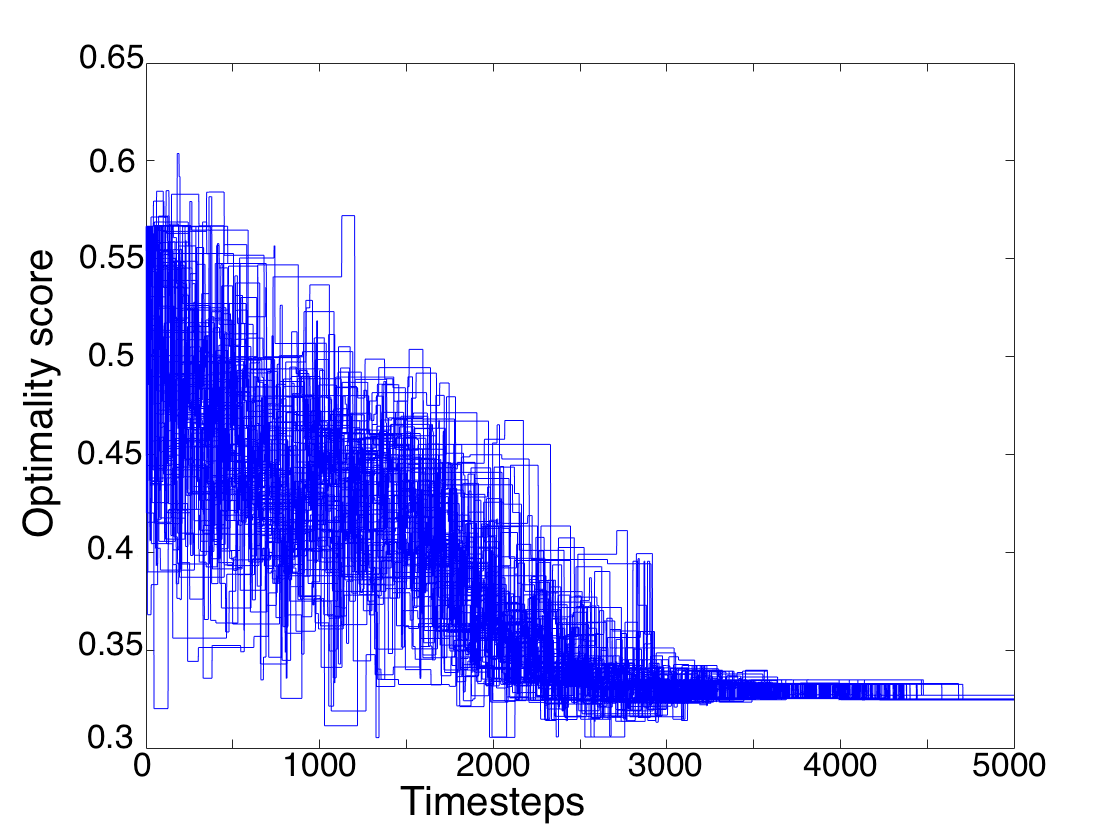

### timevarinH4.png

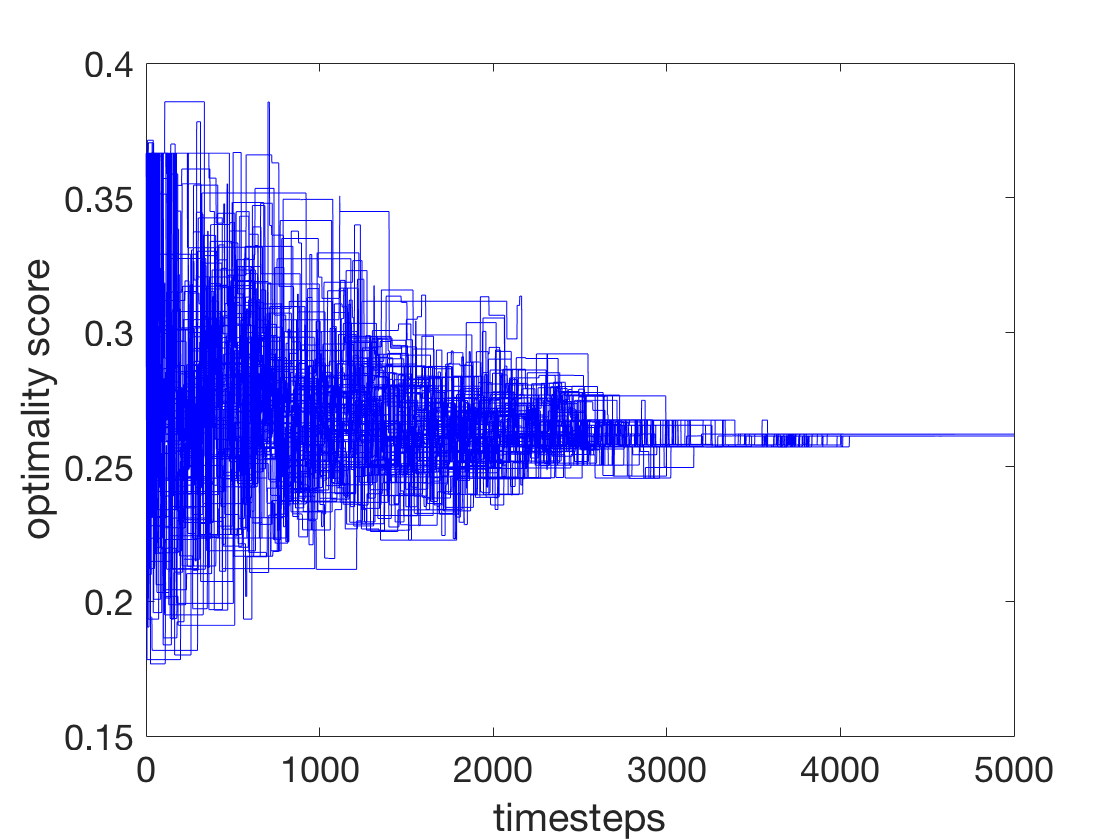
